## Supplementary information for "Conformational decoupling in acid-sensing ion channels uncovers mechanism and stoichiometry of PcTx1-mediated inhibition"

### Contents:

Supplemental Figures S1-S5
Supplemental Tables S1-S9

### Supplemental Figures

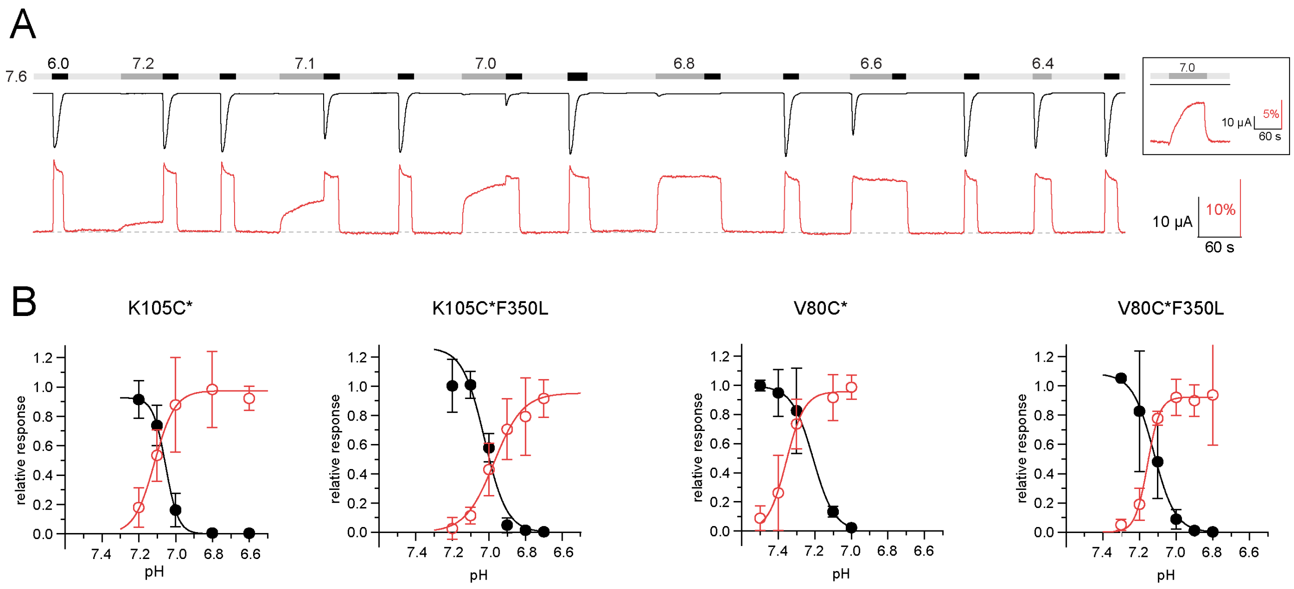


**Figure S1. VCF data of pH activation. (A)** Representative trace of a VCF recording of K105C* with the current in black and the fluorescence in red to establish steady-state desensitization (SSD) and pH-dependent fluorescence response. Inset shows pH 7.0 application without subsequent pH 5.5 activation. **(B)** SSD and fluorescence response curves for different ASIC1a variants based on recording protocols shown in (A). Data is presented as mean ± 95CI, Panel B of K105C* is adapted from (Borg, Braun et al. 2020).

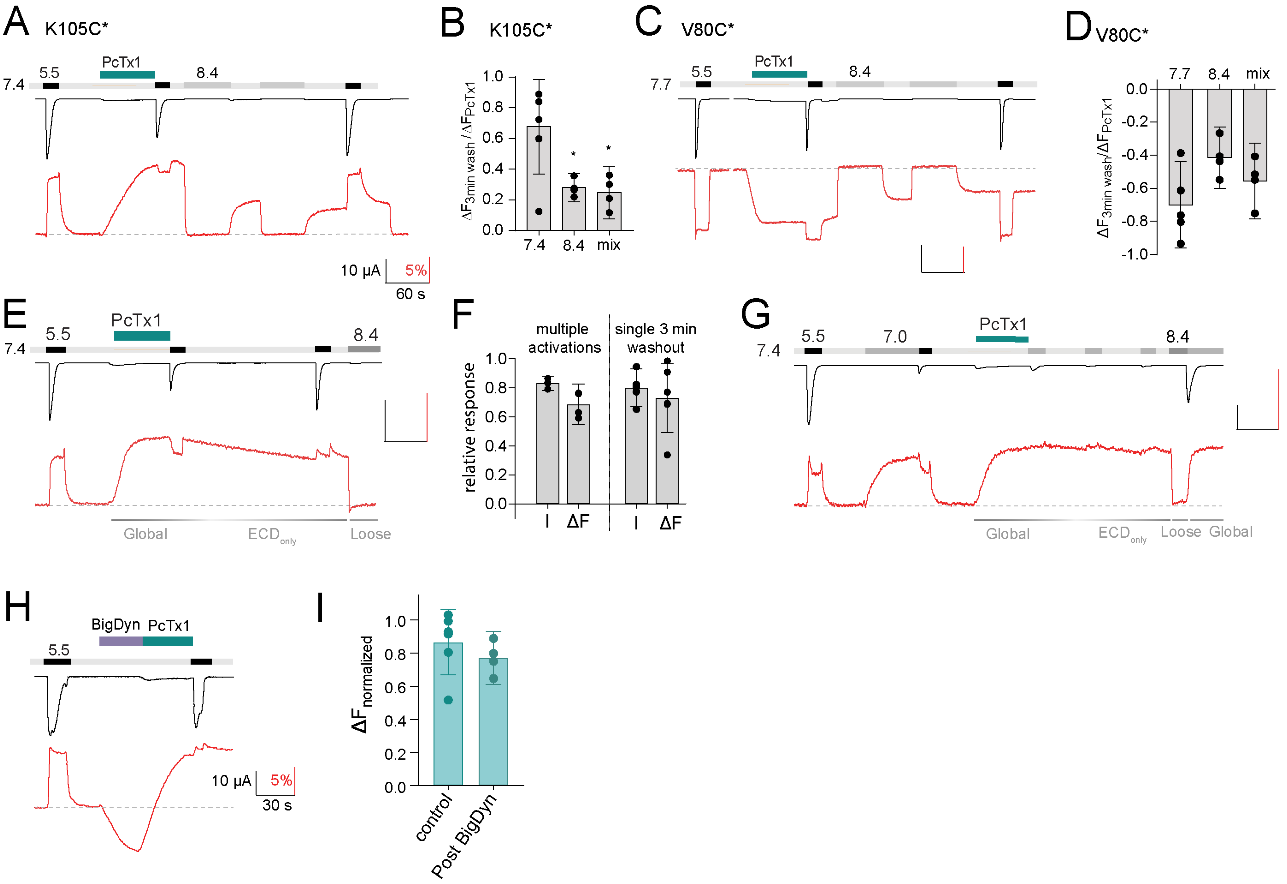


**Figure S2. VCF data of PcTx1 washout. (A)** Representative trace of a VCF recording of K105C* showing application of PcTx1 (300 nM) washed off for 3 min with alternating pHs (20 s pH 7.4, 60 s pH 8.4, 40 s pH 7.4, 60 s pH 8.4) before switching to pH 7.4 again. **(B)** Comparison of the fluorescence of K105C* after a 3 min washout of 300 nM PcTx1 using pH 7.4, 8.4 or a mix of the two as shown in the protocol in (A) and Fig. 2B. **(C)** Same as in (A) but for V80C* and running buffer 7.7 instead of 7.4. **(D)** Same as in (B) but for V80C* and base on the protocol shown in (C) and Fig. 2D). **(E)** Representative VCF trace of K105C* showing application of PcTx1 (300 nM) washout for 3 min before pH 5.5 stimulus. Corresponding binding modes are indicated below. **(F)** Comparison of the final pH 5.5-induced current (I) and fluorescence (ΔF) at pH 7.4 in recordings shown in Fig. 2A, where channels undergo three 1 min washouts at pH 7.4, each followed by pH 5.5 stimulus (left) or the protocol shown in (E) with a single 3 min washout (right). The final pH 5.5-induced current was normalized to the one at the beginning of the recording, the fluorescence was analysed right before the final pH 5.5 activation and normalized to deflection induced by PcTx1. **(G)** Representative VCF trace of K105C* showing pH 7.0 stimuli at various states of the recording. Corresponding binding modes are indicated below. **(H)** Representative trace of a VCF recording of K105C* depicting 30 s pre-conditioning with BigDyn (1 µM) and subsequent 30 s PcTx1 (300 nM) application and pH 5.5 activation. **(I)** Quantitative comparison of the fluorescence change induced by 300 nM PcTx1 at pH 7.4 in the apo state (control) and after BigDyn (1 µM) application, normalized to the signal induced by pH 5.5. Data is presented as mean ± 95CI, ordinary analysis of variance (ANOVA)

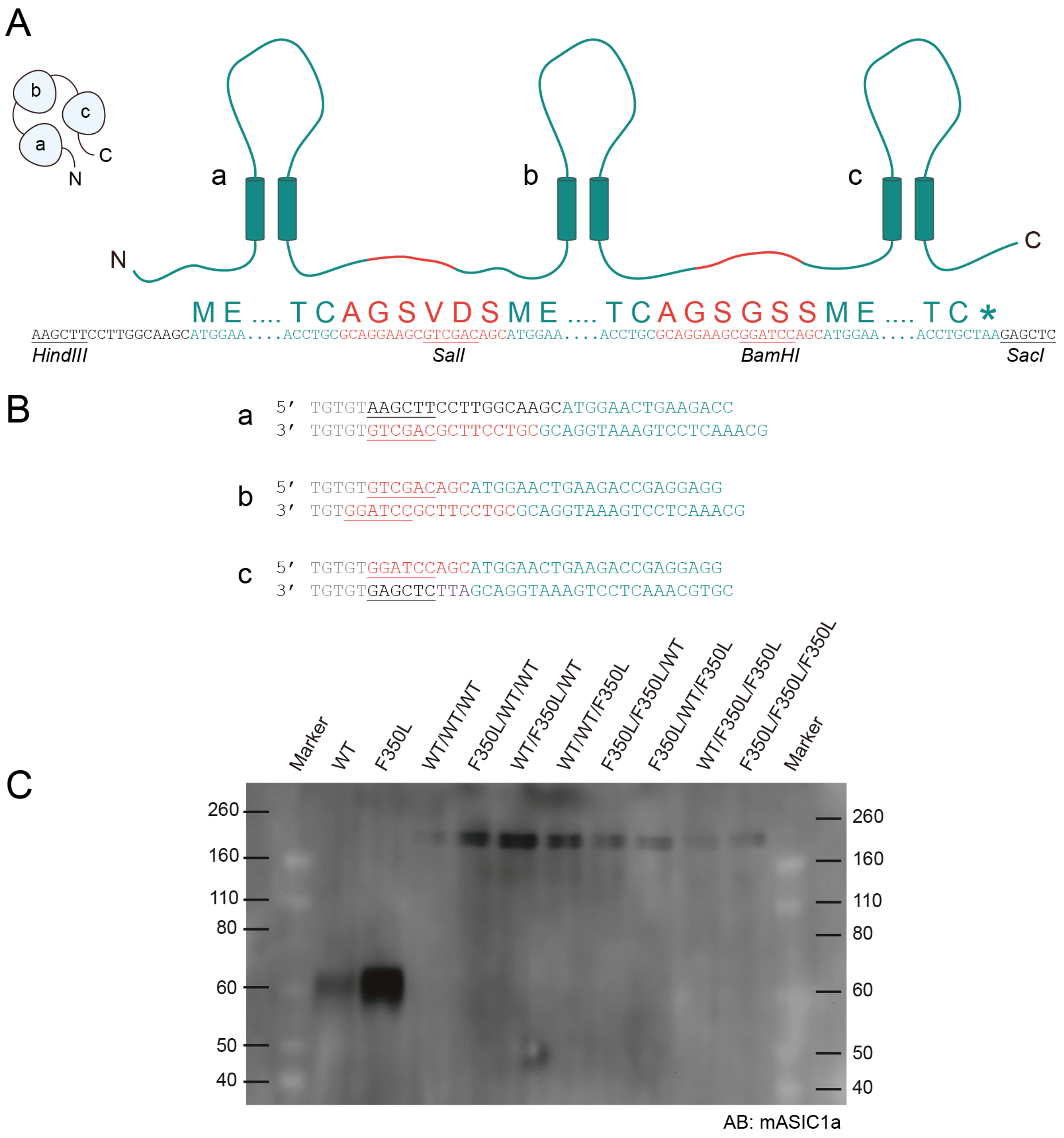


**Figure S3.** Concatemer design and validation. (A) Schematic representation of the concatemer design showing mASIC1a sequence in teal, linkers that connect subunits a, b, and c in red, with restriction sites underlined. (B) Primer pairs used to generate inserts a-c for ligation into the concatemer constructs. Nucleotide spacers and untranslated sequences are shown in grey, restriction sites are shown in red, peptide linker are shown in black and mASIC1a sequence is in teal. (C) Western blot of surface-purified membrane protein from oocytes expressing monomeric or concatemeric ASIC1a constructs using a mASIC1a specific antibody. Bands for monomeric ASIC1a WT and F350L subunits appear at ~60 kDa while the bands for concatemeric ASIC1a constructs appear at ~180 kDa, corresponding to three covalently linked ASIC1a subunits. Panels (A, B) adapted from (Lynagh, Flood et al. 2017).

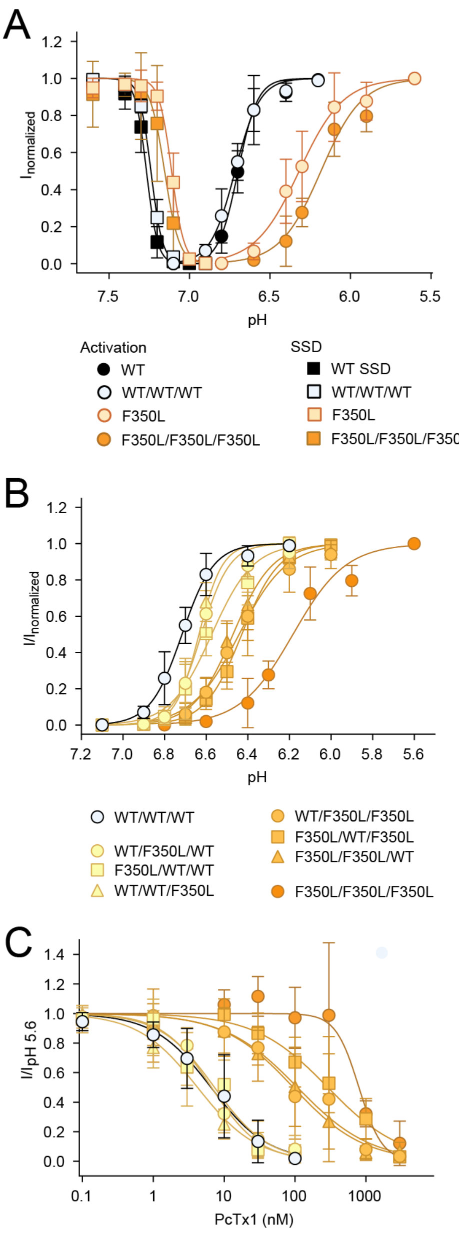


**Figure S4. Sensitivity of concatemers to pH and PcTx1. (A)** Activation and SSD curves for trimeric and concatemeric WT and F350L channels in comparison. **(B)** Activation curves for all concatemeric variants showing that concatemers with the same number of F350L-bearing subunits cluster around similar pH sensitivities. **(C)** Same as in (B), but for concentration-dependent PcTx1 inhibition.

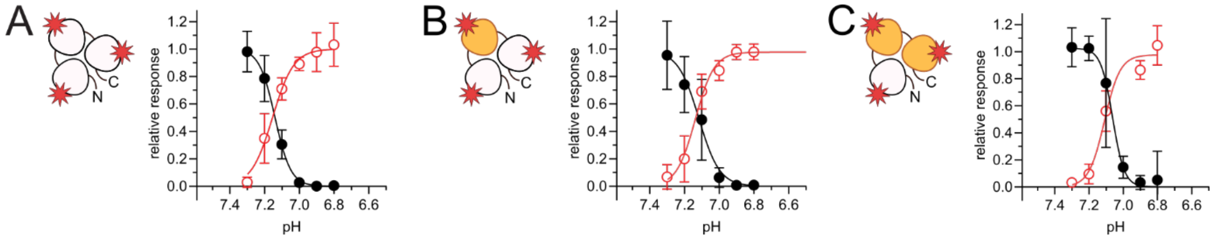


**Figure S5. VCF data of pH-dependent desensitisation and fluorescence response of concatemers. (A)** SSD and fluorescence response curves for the WT*/WT*/WT* concatemer. **(B)** Same as in (A), but for WT*/F350L*/WT* **(C)** Same as in (A), but for WT*/F350L*/F350L*.

### Supplemental Tables

**Table S1.** Effect of PcTx1 on activation and SDD of WT and F350L mASIC1a. Data is reported as mean with 95CI in brackets.

| | WT | | | F350L | | |
|---|---|---|---|---|---|---|
| PcTx1 (nM) | $pH_{50}$ | Hill | n | $pH_{50}$ | Hill | n |
| Activation | | | | | | |
| 0 | 6.68 (6.64, 6.72) | 6.16 (2.74, 9.58) | 18 | 6.26 (6.17, 6.35) | 3.61 (2.75, 4.47) | 12 |
| 30 | 7.21 (7.12, 7.30) | 2.72 (2.08, 3.36) | 7 | 6.20 (6.06, 6.36) | 3.76 (3.02, 4.53) | 7 |
| Steady-state desensitization | | | | | | |
| 0 | 7.28 (7.22, 7.33) | 14.31 (6.20, 22.43) | 6 | 7.12 (7.09, 7.16) | 19.51 (12.79, 26.22) | 5 |
| 30 | 7.68 (7.63, 7.72) | 17.15 (7.04, 27.27) | 9 | 7.28 (7.21, 7.36) | 18.78 (5.36, 32.20) | 7 |

**Table S2.** SSD and pH-dependent changes in fluorescence for labelled mASIC1a constructs. Data is reported as mean with 95CI in brackets.

| | $pH_{50}$ | Hill | n |
|---|---|---|---|
| **Steady-state desensitization** | | | |
| K105C* | 7.06 (7.02, 7.09) | 13.96 (10.88, 17.03) | 4 |
| WT*/WT*/WT* | 7.14 (7.12, 7.16) | 11.56 (7.87, 15.25) | 6 |
| WT*/F350L*/WT* | 7.13 (7.07, 7.19) | 7.84 (4.73, 10.95) | 4 |
| WT*/F350L*/F350L* | 7.06 (7.02, 7.10) | 15.04 (6.90, 23.18) | 4 |
| K105C*F350L | 6.99 (6.96, 7.03) | 11.66 (7.34, 15.98) | 4 |
| V80C* | 7.17 (7.13, 7.21) | 11.95 (5.34, 18.57) | 4 |
| V80C*F350L | 7.31 (6.72, 7.89) | 8.56 (0.03, 17.15) | 4 |
| **Fluorescence signal** | | | |
| K105C* | 7.11 (7.15, 7.06) | 7.24 (5.40, 9.07) | 5 |
| WT*/WT*/WT* | 7.15 (7.18, 7.12) | 8.07 (5.54, 10.59) | 6 |
| WT*/F350L*/WT* | 7.14 (7.11, 7.16) | 9.09 (5.06, 13.12) | 5 |
| WT*/F350L*/F350L* | 7.07 (7.15, 7.00) | 7.72 (3.70, 11.74) | 4 |
| K105C*F350L | 6.95 (6.86, 7.52) | 6.69 (3.91, 9.47) | 6 |
| V80C* | 7.36 (7.30, 7.42) | 9.41 (6.62, 12.21) | 4 |
| V80C*F350L | 7.16 (7.14, 7.18) | 13.87 (8.68, 19.06) | 5 |

**Table S3.** Resulting fluorescence change upon application of 300 nM PcTx1 for mASIC1a labelled constructs. Data is reported as mean with 95CI in brackets.

| **Construct** | **Application pH** | $\Delta F_{PcTx1}/\Delta F_{pH\ 5.5}$ | **n** |
|---|---|---|---|
| K105C* | 7.4 | 1.13 (1.09, 1.18) | 14 |
| WT*/WT*/WT* | 7.4 | 1.25 (1.01, 1.50) | 6 |
| WT*/F350L*/WT* | 7.4 | 0.99 (0.86, 1.12) | 8 |
| WT*/F350L*/F350L* | 7.4 | 0.70 (0.46, 0.95) | 5 |
| K105C*/F350L | 7.4 | 0.019 (0.001, 0,04) | 6 |
| K105C*/F350L | 7.3 | 0.464 (0.24, 0.68) | 5 |
| V80C* | 7.7 | -0.86 (-0.90, -0.81) | 13 |
| V80C*/F350L | 7.4 | -0.70 (-0.87, -0.53) | 8 |

**Table S4.** Change in fluorescence after a 3 min washout of 300 nM PcTx1. Data reported as mean with 95CI in brackets.

| Construct | Wash pH (3 min) | $\Delta F_{3min}/\Delta F_{PcTx1}$ | n |
|---|---|---|---|
| K105C* | 7.4 | 0.68 (0.37, 0.98) | 6 |
| | 8.4 | 0.28 (0.19, 0.37) | 4 |
| | mix | 0.25 (0.08, 0.42) | 4 |
| WT*/WT*/WT* | 7.4 | 0.83 (0.78, 0.88) | 6 |
| WT*/F350L*/WT* | 7.4 | 0.49 (0.39, 0.60) | 8 |
| WT*/F350L*/F350L* | 7.4 | 0.15 (0.33, 0.04) | 5 |
| K105C*/F350L | 7.4 | 0.03 (-0.01, 0.08 ) | 5 |
| V80C* | 7.7 | -0.70 (-0.96, -0.44) | 5 |
| | 8.4 | -0.41 (-0.60, -0.23) | 4 |
| | mix | -0.56 (-0.79, -0.33) | 4 |
| V80C*/F350L | 7.4 | -0.059 (-0.13, 0.01) | 6 |

**Table S5.** Comparison of current and fluorescence in experiments where 300 nM PcTx1 were applied at pH 7.4. The channels then undergo three 1 min washouts each followed by pH 5.5 stimulus, or a single 3min washout with pH 7.4 followed by a pH 5.5 stimulus. The final pH 5.5 current was normalized to the one at the beginning of the recording, the fluorescence was analyzed at pH 7.4 right before the final pH 5.5 activation and normalized to deflection induced by PcTx1. Data is reported as mean with 95CI in brackets.

| Treatment | $I_{7.4}/I_{PcTx1}$ | $\Delta F_{7.4}/\Delta F_{PcTx1}$ | n |
|---|---|---|---|
| 3 min washout | 0.79 (0.69, 0.89) | 0.73 (0.49, 0.96) | 6 |
| Multiple activations | 0.83 (0.78, 0.88) | 0.68 (0.54, 0.82) | 4 |

**Table S6.** Fluorescence after PcTx1 applications at pH 8.0 followed by different pH regimes (pH 5.5, 7.4 or 8.0) and a washout with pH 7.4. Fluorescence is reported 1 min into the final pH 7.4 application. Data reported as mean with 95CI in brackets.

| Treatment | $\Delta F_{7.4}/\Delta F_{5.5}$ | n |
|---|---|---|
| PcTx1 at pH 8.0 | -0.08 (-0,14, -0.02) | 7 |
| 7.4 after 5.5 | 0.95 (0.81, 1.08) | 6 |
| 7.4 after 7.4 | 0.69 (0.60, 0.78) | 4 |
| 7.4 after 8.0 | 0.11 (0.01, 0.23) | 4 |

**Table S7.** Resulting fluorescence change upon peptide application K105C* mASIC1a normalized to the fluorescence response elicited by pH 5.5 application. BigDyn was applied at a concentration of 1 µM and PcTx1 was applied at a concentration of 300 nM. The fluorescence change was monitored 1 min after application for BigDyn and 30–60 s after application for PcTx1. Data is reported as mean with 95CI in brackets.

| Treatment | $\Delta F_{peptide}/\Delta F_{pH\ 5.5}$ | n |
|---|---|---|
| BigDyn (Control) | -0.84 (-1.05, -0.64) | 6 |
| BigDyn post PcTx1 | 0.50 (0.18, 0.83) | 6 |
| PcTx1 (Control) | 0.86 (0.67, 1.06) | 6 |
| PcTx1 post BigDyn | 0.77 (0.61, 0.93) | 4 |
| PcTx1 (Control) | 1.20 (1.05, 1.34) | 6 |
| PcTx1 post PcTx1 | 1.17 (1.06, 1.28) | 6 |

**Table S8.** Summary of PcTx1 inhibition of mASIC1a constructs. Data is reported as mean with 95CI in brackets.

| | $IC_{50}$ | Hill | |
|---|---|---|---|
| **Construct** | **Mean (nM)** | **Mean** | **n** |
| WT | 0.6 (0.4, 0.9) | 0.9 (0.7, 1.2) | 9-14 |
| F350L | 977.2 (-, -) | 11.2 (- , -) | 4-7 |
| WT/WT/WT | 6.5 (5.2, 9.0) | 1.1 (0.9, 1.3) | 6-9 |
| F350L/WT/WT | 7.3 (5.9, 9.0) | 1.1 (0.9, 1.4) | 5-6 |
| WT/F350L/WT | 6.6 (5.7, 7.7) | 1.3 (1.0, 1.5) | 5-7 |
| WT/WT/F350L | 4.2 (3.2, 4.8) | 1.0 (0.9, 1.2) | 5-7 |
| F350L/F350L/WT | 94.6 (74.4, 120.1) | 0.9 (0.7, 1.1) | 4-6 |
| F350L/WT/F350L | 289.9 (218.1, 387.2) | 0.8 (0.7, 1.1) | 6-11 |
| WT/F350L/F350L | 108.0 (834, 141.8) | 0.8 (0.7, 1.0) | 4-7 |
| F350L/F350L/F350L | 797.1 (654.1, 962.4) | 2.6 (1.7, -) | 4-6 |

**Table S9.** pH sensitivity of activation and SSD for mASIC1a constructs. Data reported as mean with 95CI in brackets.

| Construct | $pH_{50}$ | Hill | n |
|---|---|---|---|
| **Activation** | | | |
| WT | 6.69 (6.65, 6.73) | 7.70 (5.88, 9.53) | 10 |
| WT/WT/WT | 6.72 (6.69, 6.75) | 6.31 (4.92, 7.71) | 10 |
| F350L/WT/WT | 6.59 (6.53, 6.64) | 5.80 (2.87, 8.74) | 8 |
| WT/F350L/WT | 6.62 (6.58, 6.65) | 8.82 (6.40, 11.24) | 13 |
| WT/WT/F350L | 6.62 (6.58, 6.66) | 7.72 (4.79, 10.66) | 10 |
| F350L/F350L/WT | 6.47 (6.44, 6.50) | 6.29 (4.26, 8.32) | 13 |
| F350L/WT/F350L | 6.43 (6.39, 6.46) | 5.34 (4.75, 5.92) | 8 |
| WT/F350L/F350L | 6.44 (6.37, 6.50) | 6.96 (4.15, 9.77) | 10 |
| F350L/F350L/F350L | 6.18 (6.11, 6.26) | 3.86 (2.56, 5.17) | 7 |
| F350L | 6.31 (6.18, 6.43) | 4.09 (2.87, 5.30) | 7 |
| **Steady-state desensitisation** | | | |
| WT | 7.27 (7.25, 7.30) | 14.15 (10.20,18.09) | 11 |
| WT/WT/WT | 7.24 (7.22, 7.25) | 12.11 (10.72, 13.50) | 5 |
| WT/F350L/WT | 7.24 (7.19, 7.29) | 13.79 (8.62, 18.97) | 5 |
| WT/F350L/F350L | 7.17 (7.14, 7.20) | 9.09 (5.77, 12.42) | 6 |
| F350L/F350L/F350L | 7.16 (7.11, 7.21) | 12.45 (4.60, 20.29) | 4 |
| F350L | 7.11 (7.09, 7.14) | 12.36 (8.96, 15.76) | 7 |
